# Emotional information facilitates or disrupts memory integration through distinct hippocampal processes of reactivation and connectivity

**DOI:** 10.1101/2023.04.25.538111

**Authors:** Yannan Zhu, Wei Liu, Nils Kohn, Guillén Fernández

## Abstract

Emotion has a significant impact on how related experiences are organized into integrated memories. However, the neurobiological mechanisms of how emotion modulates memory integration for related information with different valences remain unclear. In this between-subject functional magnetic resonance imaging (fMRI) study, we investigated different emotional modulations of memory integration by manipulating the valence of stimuli used in an associative memory paradigm. Three groups of participants were tested: one group integrated emotional (i.e., negative) information with neutral information, one group integrated two emotional pieces of information, and one control group integrated two neutral pieces of information. Behaviorally, emotional information facilitated its integration with neutral information but interfered with the other emotional information. Neurally, the emotion-induced facilitation effect, occurring on memory integration of neutral and emotional information, was associated with increased trial-specific reactivation in the hippocampus during both encoding and retrieval. This facilitated integration was also supported by strengthened hippocampal connectivity with the amygdala, as well as a set of neocortical areas related to emotion regulation and the default mode network (DMN). In contrast, the emotion-induced interference effect, occurring on memory integration of two emotional pieces of information, was associated with impaired hippocampal trial-specific reactivation during retrieval that appeared to offset the facilitating effect of increased reactivation during encoding. Similar but relatively weak hippocampal connectivity was found underlying this interfered integration. Taken together, emotional information facilitates memory integration with neutral information, while disrupting the integration with other emotional information, through distinct dynamical processes of hippocampal trial-specific reactivation and connectivity.

## Introduction

Our episodic memory is not a static repository of experiences, but a dynamically and constantly updating system enabling future use in an ever-changing environment (Ebbinghaus, 1885; Nader et al., 2000; Tulving, 1983). Memories of related experiences can be integrated into a highly adaptive network of overlapping representations (Eichenbaum, 2000; Shohamy and Wagner, 2008; Tulving, 1995). This memory integration mechanism makes interactions between existing knowledge and new experiences possible, contributing to a set of advanced cognitive functions, such as generalization (McClelland et al., 1995; Shepard, 1987), inference (Bunsey and Eichenbaum, 1996; Zeithamova, Schlichting, et al., 2012), and schema (Bartlett, 1932; Piaget, 2002). Through memory integration, emotional arousal not only strengthens memory for an emotional event (LaBar and Cabeza, 2006; LeDoux, 1994; Schacter, 1999), but also has a broader impact on related, other events. Despite considerable efforts, a general consensus on the emotional modulation of related events has not been reached, as both facilitation and interference effects reported in divergent studies (Bravo-Rivera and Sotres-Bayon, 2020; Li et al., 2008; Mather and Sutherland, 2011; Wimmer and Shohamy, 2012; Zhu et al., 2022). In this study, we manipulated the emotional composition of to-be-integrated stimuli to elucidate the circumstances under which facilitation and interference effects occur, as well as their potentially distinct neural mechanisms.

In our daily life, an emotional experience can modulate memories of specific events that share common contents with the emotional event even though they occur separately in time. Such episode-unique emotional learning can be tested with a trial-specific associative memory paradigm, accommodated by the memory integration mechanism (Bunsey and Eichenbaum, 1996; Preston et al., 2004). Given the view of interactive relationships among integrated memories (Bartlett, 1932; Schlichting and Frankland, 2017; Schlichting and Preston, 2015), it is conceivable that the emotional modulation of memory integration not only depends on emotional information itself but also the valence of other related information in the integration. One prominent view, “facilitation theory”, has proposed that emotion facilitates memory integration (Holmes et al., 2022; Shohamy and Daw, 2015). Recent studies have shown that an emotionally salient experience enhances memory for related neutral events, indicating a tighter memory integration with strengthened associations among these pieces of information (Wang and Kahnt, 2021; Wimmer and Shohamy, 2012; Wong et al., 2019; Zhu et al., 2022). However, an “interference theory” suggests that emotion disrupts memory integration (Bravo-Rivera and Sotres-Bayon, 2020; Mather, 2007). Studies supporting this view provide evidence of a mutual inhibitory effect, where equally prioritized emotional memories compete with each other and are overall suppressed (Mather and Sutherland, 2011; Morelli and Burton, 2009). Both facilitation and interference theories have gained supports from experimental evidence. However, clear evidence of when emotional information facilitates and when it disrupts memory integration is still needed.

Memory integration is thought to firstly form during encoding, and subsequently update during retrieval (de Sousa et al., 2021; Holmes et al., 2022). Human neuroimaging studies provide compelling evidence of an integrative encoding mechanism, also known as online integration (Kuhl et al., 2010; Shohamy and Wagner, 2008; van Kesteren et al., 2016). It proposes that the neural pattern of initial memory can be reactivated during the encoding of a new event with their overlapping representations as a bridge. Our memories become malleable to modification when reactivated, and thus can be integrated into a linked mnemonic network (Przybyslawski and Sara, 1997; Schwabe et al., 2014). Besides the encoding phase, a large body of evidence shows that memory retrieval also triggers the reactivation of neural ensembles engaged during encoding of the event (Liu et al., 2020; Nyberg et al., 2000; Tayler et al., 2013; Tulving and Thomson, 1973). The interactive influence between reactivation of integrated memories during retrieval could modify their initially formed integration (Bauml and Samenieh, 2010; Carneiro et al., 2021; Roediger and Abel, 2022). However, there is a gap in research on the various emotional modulations of memory integration, especially regarding how this process evolves from encoding to retrieval.

The hippocampus has been well recognized to play a critical role in integrating discrete experiences into a cohesive memory network through pattern completion (Eichenbaum, 2000; McClelland et al., 1995; Treves and Rolls, 1994), and might lie at the heart of differential emotional memory integration effects. Reactivation of hippocampal representations during the encoding of new events and retrieval supports the flexible integration of related experiences (Biderman et al., 2020; Kuhl et al., 2010; Tarder-Stoll et al., 2021; Wimmer and Buchel, 2021). Besides reactivation, hippocampal coordinated interactions with neocortical or subcortical structures, especially emotion-related regions (e.g., the amygdala), are also thought to be involved in emotional memory integration (Lima Portugal et al., 2020; Remondes and Wilson, 2013; Richter-Levin and Akirav, 2000; Sutherland and McNaughton, 2000). However, it remains unclear how hippocampal reactivation and connectivity contribute to different emotional modulations of memory integration.

Here, we examined the emotion-related facilitation and interference effects for memory integration. We conducted a between-subject fMRI experiment with an associative memory paradigm to probe memory integration, in which participants learned ABC triplets with two location-item associations (i.e., AB and AC) sharing one location cue (i.e., A) (**Fig. 1A**). To investigate memory integration between emotional (i.e., negative) information and related information with different valences (i.e., neutral or negative) across participants, we formed three groups: one integrating neutral and emotional associations (i.e., Neutral-Emotional group), one integrating two emotional associations (i.e., Emotional-Emotional group), and one control group integrating two neutral associations (i.e., Neutral-Neutral group). First, we investigated how emotional information modulates integrated memory performance across the three groups. Then, we conducted pattern similarity and connectivity analyses to further investigate how hippocampal reactivation and connectivtity during the encoding and retrieval phases may account for the potentially different emotional modulations.

**Figure 1.**
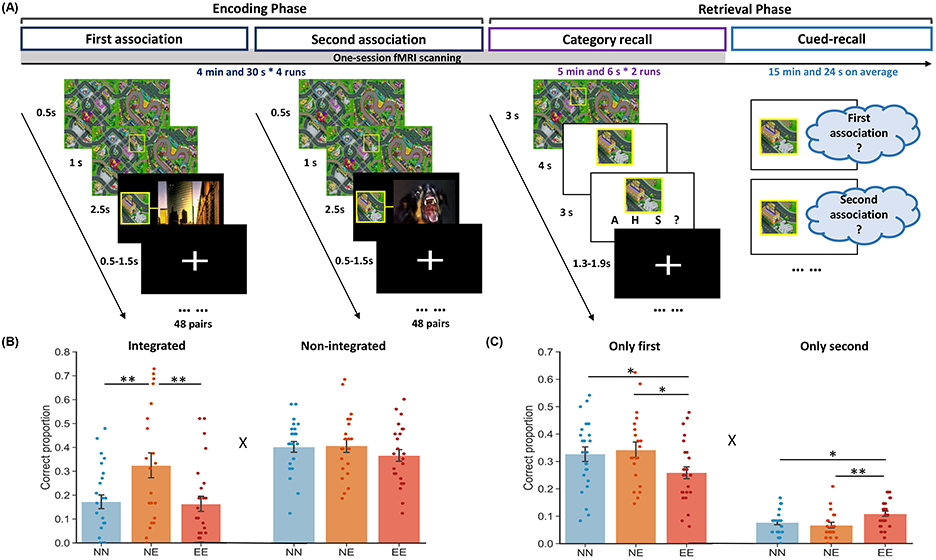
Experimental design and cued-recall behavioral performance. (A) The experiment consisted of an encoding phase and an immediate retrieval phase including two memory tests. During the encoding phase, participants learned 48 first associations and then 48 overlapping second associations. During the retrieval phase, participants performed a category recall test only for the first associations and subsequently performed a cued-recall test for both the first and second associations. (B) Bar graphs depict average correct proportions for triplets with both associations remembered (i.e., integrated memory), and triplets with only one (either first or second) association remembered (i.e., non-integrated memory) in the three groups separately. (C) Bar graphs depict average proportions for triplets with only the first association remembered (i.e., only first associative memory), and triplets with only the second association remembered (i.e., only second associative memory) in the three groups separately. Error bars represent the standard error of the mean. Dots represent data from each participant. “X” indicates (marginally) significant interaction (p < 0.085). Notes: *p < 0.05; **p < 0.01; two-tailed tests.

## Materials and methods

### Participants

A total of 76 young, healthy, native Dutch-speaking participants were recruited from the Radboud University Subject Pool (48 females; aged 18-33 years; mean age ± s.d., 23.4 ± 3.20 years). Data from six participants were excluded from the following analyses due to failure to complete all tasks (n = 5), or failure to understand or follow the task instructions (n = 1). Participants were randomly assigned into one of the three groups, one group in which participants encountered two neutral items associated with one location for each triplet (Neutral-Neutral group referred to as “NN group” in the text; n = 25, 14 females), one group in which participants encountered a location with a neutral item followed by an emotional item for each triplet (Neutral-Emotional group referred to as “NE group”; n = 21, 16 females), and the third group in which participants encountered two emotional items associated with each location (Emotional-Emotional group referred to as “EE group”; n = 24, 16 females). Two additional participants in the EE group were excluded from analyses of the category recall test due to falling asleep during scanning. The sample size was determined based on prior studies using associative memory paradigm (Schlichting et al., 2021; Schlichting et al., 2015). The experiment was approved by and conducted following the requirements of the local ethics committee (CMO2014/288, Commissie Mensgebonden Onderzoek, Region Arnhem-Nijmegen, The Netherlands), and the declaration of Helsinki, including the requirement of written informed consent from each participant before the beginning of the experiment.

### Materials

Stimuli included 48 locations on two cartoon maps and 192 pictures. Specific locations (e.g., hospital, restaurant, and square), as memory cues, were captured from two cartoon maps with 24 locations in each map (Liu et al., 2020, 2021). The distinctive pictures (i.e., 96 negative and 96 neutral) were selected from the International Affective Picture System (Lang et al., 1997). Each picture belongs to one of the three categories: animal (e.g., barking dog), human (e.g., reading girl), and scene (e.g., modern city). Category information was used for the category recall test under scanning to enable simple button press responses. Each location cue (i.e., A) was paired with two items (i.e., B and C) to create 48 ABC triplets. In the NN group, both items B and C were neutral pictures. In the NE group, items B were neutral and items C were negative pictures. In the EE group, both items B and C were negative pictures. Items B and C were selected from different categories and counterbalanced for individual participants in each group.

### Experimental procedures

Participants were instructed to learn 48 location-item pairs (i.e., AB) as first associations, and then the same 48 locations paired with new items (i.e., AC) as second associations during a blocked design associative encoding phase. Thereafter, during the retrieval phase, participants completed a category recall test. The associative encoding and category recall test were performed during a 30-min fMRI scanning session. After an approximately 20-min delay, a self-paced cued-recall test was performed outside the scanner (**Fig. 1A**).

#### Associative encoding

This phase included four 4 min and 30 s runs. In each run, participants learned 12 first associations, and then 12 corresponding second associations twice (i.e., AB, AC, AB, AC). They were instructed to vividly imagine each location in relation to its paired item to aid their memory. During each trial, the entire map was initially presented for 0.5 s, followed by a yellow frame appearing on the map to highlight a specific location cue for 1 s, and then the location cue and its paired item were presented side-by-side together for 2.5 s. Trials were interleaved by a fixation cross with an inter-trial interval jittered from 0.5 to 1.5 s (i.e., 1 s on average with 0.25 s steps).

#### Category recall

Subsequently, participants performed an immediate category recall test for the 48 first associations. This task included two 5 min and 6 s runs with 24 trials in each run. During each trial, the entire map with a highlighted location was presented for 3 s and followed by the location cue on its own for 4 s. Participants were instructed to recall the item paired with this location cue in the FIRST association (i.e., AB) as vividly as possible, and then indicate the category of their imagined item by pressing an appropriate button from the four options (i.e., Animal, Human, Scene, and Don’t know) within 3 s. Trials were also interleaved by a fixation cross with an inter-trial interval jittered from 1.3 to 1.9 s (i.e., 1.6 s on average).

#### Cued-recall

After completing the category recall test during the scanning session, participants performed a self-paced cued-recall test with an average duration of 15 min and 24 s (i.e., range from ∼8 to ∼34 min), for both first and second associations. Each of the 48 location cues was presented in a random order across participants. Participants were instructed to recall paired item in the first and second associations separately, by typing a brief description of this item on a standard keyboard within 60s.

### Behavioral data analysis

Participants’ demographic data and memory performance were analyzed using Statistical Product and Service Solutions (SPSS, version 23.0, IBM). In the cued-recall test, participants’ description answers were evaluated by two native Dutch experimenters independently. If the answer provided enough detailed information (e.g., a little black cat) for the experimenter to identify the correct item in the association, as distinct from other items used in the experiment, it was labeled as correct. If the answer was detailed enough but allowed the experimenter to identify the item in the other association of this triplet (i.e., an answer of item in the corresponding second association when the instruction was to recall the first association, and vice versa), it was labeled as related. Otherwise, if the answer was entirely wrong (i.e., an answer of item in neither the first nor second association of this triplet) or not specific enough (e.g., a small animal), then it was labeled as incorrect. We used Cohen’s kappa coefficient (K) to measure inter-rater reliability (Altman, 1990), and found almost perfect reliability between the two experimenters’ evaluations (K = 0.96, *p* < 0.001). For the answers in which the experimenters disagreed (3.28 trials on average with a total of 96 trials per participant), they were resolved by discussion between the two experimenters or by extra evaluation from a third experimenter. The cued-recall memory performance was calculated based on the final determination. Effect sizes reported for ANOVAs are partial eta squared, referred to in the text as η^2^. 95% confidence intervals (CI) for post-hoc comparisons were also reported. Effect sizes reported for paired t-tests are Cohen’s *d*.

### Imaging acquisition

MRI data were acquired using a 3.0 T Siemens Skyra (Siemens Medical, Erlangen, Germany) with a 32-channel head coil system at the Donders Institute, Centre for Cognitive Neuroimaging in Nijmegen, the Netherlands. Functional images were collected using a multi-band echo-planar imaging (mb-EPI) sequence (slices, 66; multi-slice mode, interleaved; slice thickness, 2 mm; TR, 1000 ms; TE, 35.2 ms; flip angle, 60°; multiband accelerate factor, 6; voxel size, 2 × 2 × 2 mm; FOV, 213 × 213 mm). To correct for spatial distortions, fieldmap images were acquired (slices, 66; multi-slice mode, interleaved; slice thickness, 2 mm; TR, 500 ms; TE1, 2.80 ms; TE2, 5.26 ms; flip angle, 60°; voxel size, 2 × 2 × 2 mm; FOV, 213 × 213 mm). Structural images were acquired using a three-dimensional sagittal T1-weighted magnetization-prepared rapid gradient echo (MPRAGE) sequence (slices, 192; slice thickness, 1 mm; TR, 2300 ms; TE, 3.03 ms; flip angle, 8°; voxel size, 1 × 1 × 1 mm; FOV, 256 × 256 mm).

### Imaging preprocessing

Brain imaging data were preprocessed using fMRIPrep (version 20.0.6) (Esteban et al., 2019) and FEAT (fMRI Expert Analysis Tool, version 6.0) (Jenkinson et al., 2012). Brain images were corrected for field inhomogeneity using the fieldmaps, and realigned for head-motion correction using MCFLIRT (Jenkinson et al., 2002). No additional slice-timing correction was performed. Each participant’s functional images were then co-registered to their T1-weighted anatomical image using FLIRT with the boundary-based registration (BBR) cost-function (Greve and Fischl, 2009), and spatially normalized into a standard MNI space (FSL’s MNI152 2 mm template). Images used for the univariate general linear model (GLM) and psycho-physiological interaction (PPI) analyses were smoothed with a 6-mm FWHM Gaussian kernel, whereas no spatial smoothing was performed on images used for the pattern similarity analysis to retain voxel-wise information. We performed the automatic removal of motion artifacts using independent component analysis (ICA-AROMA) (Pruim et al., 2015) on brain data to further remove spurious noise related to motion, using non-aggressive denoising. In addition, we discarded the first 10 volumes of functional images for signal equilibrium and applied high-pass temporal filtering (Gaussian-weighted least-squares straight line fitting with sigma = 90.0 s) before the subsequent analyses.

### Univariate general linear model (GLM) analysis and regions of interest (ROIs) definition

To investigate the brain activity in response to subsequently remembered relative to forgotten items (i.e., subsequent memory effect, SME), we conducted voxel-wise GLMs for the encoding phase. FILM prewhitening was also applied to remove temporal autocorrelation (Woolrich et al., 2001). Trials (i.e., both first and second associations) were modeled with 4 s from the onset of each stimulus (i.e., the entire map) and convolved with FEAT’s hemodynamic response function (HRF). Three regressors of interest were included in each GLM based on individual cued-recall memory performance: 1) correct responses; 2) related responses; 3) incorrect, missing, or ‘don’t know’ responses. To account for potential artifacts of movement, the motion parameters produced during realignment and stick functions (i.e., frame displacement that exceeded a threshold of 2 mm) were also included as additional regressors in each GLM. Contrast images of remembered (i.e., correct responses) relative to forgotten (i.e., incorrect, missing, or ‘don’t know’ responses) condition were generated at the individual-subject level. Related responses were not included in this contrast, as they were very few and might be caused by wrong order (i.e., first or second) memory. These resultant images were then entered into the group-level analysis, and corrected for multiple comparisons using cluster-mass thresholding within FEAT (voxel-wise z > 3.1, cluster-level *p* < 0.05 FWER corrected; *Fig. S1A*).

To investigate the brain activity in response to correctly recalled relative to forgotten items (i.e., successful retrieval effect), we conducted voxel-wise GLMs for the retrieval phase (i.e., the category recall test for first associations only). Trials were modeled with 3 s from the onset of each stimulus (i.e., the entire map with highlighted location) and convolved with FEAT’s HRF. The three regressors of interest (i.e., correct responses, related responses, and incorrect, missing, or ‘don’t know’ responses) based on individual category-recall performance were included in each GLM. The following 4-s presentations of location cues and 3-s presentations of category detection were also included as two regressors of no interest. Contrast images (i.e., remembered vs. forgotten) were generated at the individual-subject level and then entered into the group-level analysis (*Fig. S1B*). All other settings (i.e., motion regressors and thresholding) were the same as the above GLM analysis of the encoding phase.

We functionally identified a bilateral hippocampal ROI by the univariate activation contrast of remembered relative to forgotten condition during encoding. The hippocampal ROI defined by the encoding contrast map was contained in the anatomical automatic labeling (AAL) template of “hippocampus” constrained by the retrieval contrast map (i.e., remembered vs. forgotten). Thus, our hippocampal ROI was engaged (i.e., activate and reactivate) in both encoding and retrieval, showing a reliable memory effect. This hippocampal ROI was used in the following pattern similarity and PPI analyses (**Fig. 2B, 3A,** and *Fig. S6A*). The posterior medial cortex (PMC), as a control region in pattern similarity analyses, was defined by a probabilistic atlas of resting-state default mode network from FIND lab at Stanford University (https://findlab.stanford.edu/functional_ROIs.html) (Chen et al., 2016; Shirer et al., 2012) (*Fig. S5A*). We also anatomically defined a left amygdala ROI based on the AAL from the WFU PickAtlas toolbox (http://fmri.wfubmc.edu/software/PickAtlas). The amygdala ROI was used in the PPI analysis (**Fig. 3B** and *Fig. S6B*).

**Figure 2.**
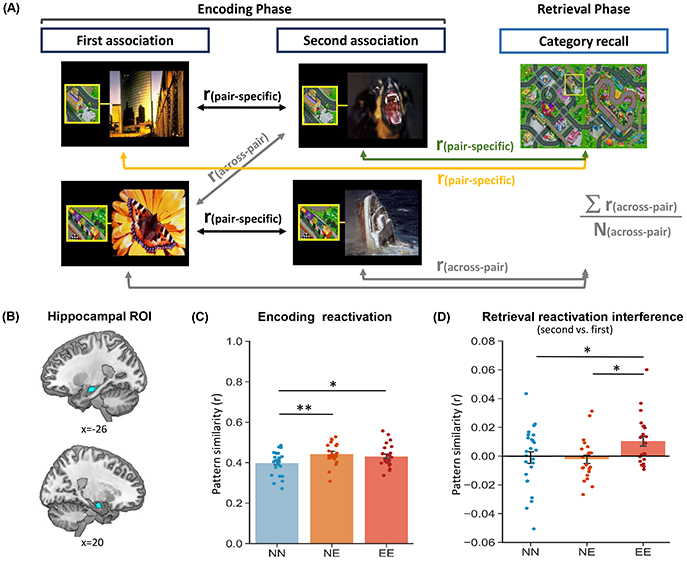
Hippocampal neural reactivation during encoding and retrieval. (A) Pattern similarity analysis approach. Several similarity measures were computed for each trial. Encoding pair-specific similarity (black) was computed by correlating each first association’s multi-voxel activity pattern with its corresponding pattern of the second association during encoding. Encoding across-pair similarity (grey) was computed by averaging all correlations between each first association’s pattern and the patterns of all other different second associations during encoding. Retrieval-first pair-specific similarity (yellow) was computed by correlating each location cue’s pattern during retrieval with its corresponding pattern of the first association during encoding. Retrieval-second pair-specific similarity (green) was computed by correlating each location cue’s pattern during retrieval with its corresponding pattern of the second association during encoding. Retrieval across-pair similarity (grey) was computed by averaging all correlations between each location cue’s pattern during retrieval and the patterns of all other different associations (i.e., both first and second associations) during encoding. These correlations from each similarity measure were then averaged across trials for each participant. (B) The functionally defined bilateral hippocampal ROI was used in the pattern similarity analysis. (C) Bar graphs depict average encoding reactivation (i.e., encoding pair-specific vs. across-pair similarity) in the three groups separately. (D) Bar graphs depict average retrieval reactivation interference (i.e., retrieval-second vs. retrieval-first reactivation) in the three groups separately. Error bars represent the standard error of the mean. Dots represent data from each participant. Notes: *p < 0.05; **p < 0.01; two-tailed tests.

**Figure 3.**
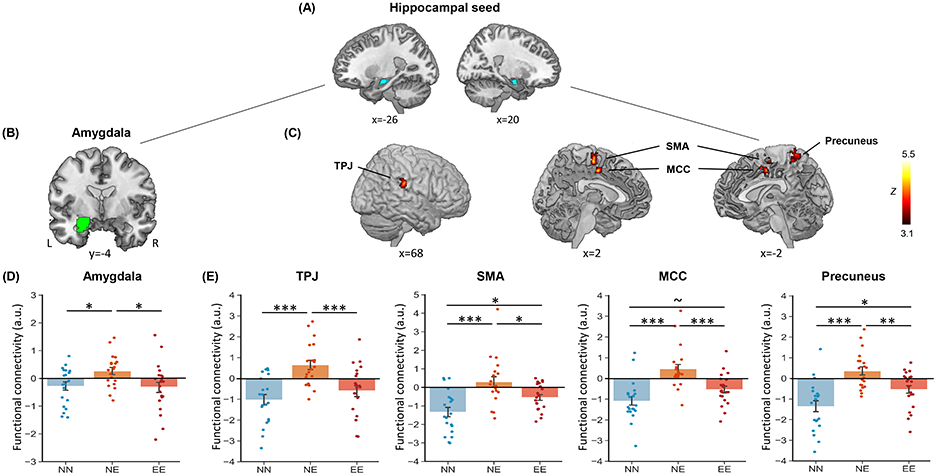
Hippocampal connectivity involved in successful memory integration during encoding. (A) The bilateral hippocampal seed was used in the task-dependent gPPI analysis. (B) The anatomically left amygdala ROI was used in this hippocampal connectivity analysis. (C) Significant clusters in the right temporoparietal junction (TPJ), bilateral supplementary motor area (SMA), bilateral middle cingulate cortex (MCC), and left precuneus, showing the main effect of Group. (D-E) Bar graphs depict average hippocampal connectivity with the amygdala, TPJ, SMA, MCC, and precuneus in the three groups separately. Error bars represent the standard error of the mean. Dots represent data from each participant. Notes: Color bar represents z values; a.u., arbitrary unit; L, left; R, right; ∼ p < 0.08; *p < 0.05; **p < 0.01; ***p < 0.001; two-tailed tests.

### Reactivation: multivariate pattern similarity analysis (MVPA)

To assess emotion-modulated effects on hippocampal reactivation during encoding and retrieval phases, we conducted an ROI-based trial-by-trial MVPA (Kriegeskorte et al., 2008; Xue et al., 2010). A separate regressor was generated for each trial with 2.5 s from the onset of each location-picture pair during the encoding phase (i.e., collapsing across two repetitions), and for each trial with 3 s from the onset of each entire map with highlighted location during the category retrieval phase, convolved with FEAT’s HRF. It resulted in four GLMs for the encoding phase and two GLMs for the retrieval phase, each with 24 regressors. Then, the *t*-values of resulting spatial activation pattern for each trial in the hippocampal ROI were extracted into a vector and z-scored. The similarity between different vectors was computed using Pearson’s correlation and Fisher-transformed before statistical analyses.

### Functional connectivity: psycho-physiological interaction (PPI) analysis

To assess emotion-modulated effects on hippocampal functional connectivity during encoding and retrieval phases, we conducted a generalized form of task-dependent PPI (gPPI) analysis (Friston et al., 1997; Harrison et al., 2017). Contrast images corresponding to the whole-brain PPI effect (i.e., second vs. first associations during encoding, or triplets with both associations remembered vs. no association remembered during category retrieval) at the individual-subject level were then submitted to a one-way ANOVA for the group-level analysis. Significant clusters were determined using the same criterion as the above univariate GLMs. Mean *t*-values extracted from the resultant significant clusters were submitted into separate post-hoc comparison tests, and visualized in bar graphs to show specific group differences. In line with our prior hypothesis, we also extracted mean *t*-values of hippocampal connectivity data from the amygdala ROI and then submitted them into the one-way ANOVA.

### Data and code availability

The research data and scripts of this study were uploaded to the Donders repository (https://data.donders.ru.nl/) and are publicly available upon publication. The project was named “Emotional Modulation of the Interaction Between Related Memories with Functional MRI” in the repository (https://doi.org/10.34973/esn0-yf75).

## Results

### Behavioral performance: Emotional information facilitates integrated memory with related neutral information, but interferes with other emotional information

Based on the cued-recall memory performance, we calculated the integrated memory performance as the proportion of triplets with both associations remembered correctly. We also calculated the non-integrated memory performance for associations that failed to integrate with each other, as the proportion of triplets with only one (i.e., either first or second) association remembered correctly. The latter was further separated into the proportions of triplets with only the first association remembered correctly, and triplets with only the second association remembered correctly, to measure only first and only second associative memory performances.

Firstly, to investigate different emotion-modulated effects on integrated memory compared to non-integrated memory across the three groups, we conducted a 2 (Memory: integrated vs. non-integrated) by 3 (Group: Neutral-Neutral vs. Neutral-Emotional vs. Emotional-Emotional) repeated-measures ANCOVA with gender as a covariate of no interest, given that gender was not balanced across groups. This analysis revealed a significant main effect of Group (F_(2,66)_ = 5.74, *p* = 0.005, partial η^2^ = 0.15) and a trend for Memory-by-Group interaction effect (F_(2,66)_ = 2.60, *p* = 0.082, partial η^2^ = 0.07), but no main effect of Memory (F_(1,66)_ = 0.47, *p* = 0.495, partial η^2^ = 0.01; **Fig. 1B**). Post-hoc comparisons revealed better integrated memory in the Neutral-Emotional (i.e., NE) group than Neutral-Neutral (i.e., NN; *p* = 0.006, 95%CI = [0.047, 0.273]) and Emotional-Emotional (i.e., EE; *p* = 0.005, 95%CI = [0.052, 0.278]) groups, but no significant difference in non-integrated memory across the three groups (all *p* > 0.290). This result indicates that emotional information selectively facilitates its integrated memory with related neutral information in the NE group.

It is worth noting that this emotion-related facilitation effect for integrated memory was not found in the EE (vs. NN) group (*p* = 0.928), although the EE group contained emotional information as well. Thus, to further investigate whether a different emotional modulation especially occurs in the EE group, we conducted a 2 (Memory: only first vs. only second) by 3 (group: NN vs. NE vs. EE) repeated-measures ANCOVA also with gender as a covariate of no interest. This analysis revealed a trending main effect of Memory (F_(1,66)_ = 3.18, *p* = 0.079, partial η^2^ = 0.05) and a significant Memory-by-Group interaction effect (F_(2,66)_ = 5.46, *p* = 0.006, partial η^2^ = 0.14), but no main effect of Group (F_(2,66)_ = 0.74, *p* = 0.484, partial η = 0.02; **Fig. 1C**). Post-hoc comparisons revealed better second associative memory in the EE group than NN (*p* = 0.011, 95%CI = [0.008, 0.061]) and NE (*p* = 0.005, 95%CI = [0.012, 0.067]) groups, but in contrast, worse first associative memory in the EE group than NN (*p* = 0.042, 95%CI = [−0.143, −0.003]) and NE (*p* = 0.036, 95%CI = [−0.152, −0.005]) groups. A similar repeated-measures ANCOVA for category-recall memory performance also revealed the Memory-by-Group interaction effect (F_(2,64)_ = 7.35, *p* = 0.001, partial ^2^ = 0.19), with better non-target second memory (*p* = 0.005, 95%CI = [0.024, 0.132]) but worse target first memory (*p* < 0.001, 95%CI = [−0.306, −0.098]) in the EE than NE group (*see Fig. S2 for details*). These results indicate that emotional information disrupts its integrated memory with the other related emotional information, perhaps due to a competition between two overlapping emotional associations in the EE group.

These behavioral results collectively demonstrate that emotional stimuli can facilitate the integration of neutral and emotional information into episodic memoy, while also disrupting the integration of two emotional pieces of information.

### Pattern similarity: The emotion-related facilitation and interference effects are separately associated with increased and impaired hippocampal reactivation during encoding and retrieval

Next, to investigate whether and how hippocampal reactivation during encoding and retrieval phases contributes to the observed emotion-related facilitation and interference effects, we conducted a trial-by-trial MVPA on the hippocampal ROI (**Fig. 2B**). The encoding reactivation characterizes neural reactivation of learning activity for first association cued by the overlapping location during the encoding of its corresponding second association. It was measured as encoding pair-specific relative to across-pair similarity to capture the trial-specific reactivation (or reinstatement) effect (**Fig. 2A**). The retrieval reactivation, including retrieval-first reactivation and retrieval-second reactivation, characterizes neural reactivation of encoding activity for first/second association cued by the location during the category retrieval. The retrieval-first/second reactivation was also measured as retrieval-first/retrieval-second pair-specific relative to retrieval across-pair similarity (**Fig. 2A**). Consistent with previous studies (Jonker et al., 2018; Wimmer and Shohamy, 2012; Zeithamova, Dominick, et al., 2012), we found that both encoding reactivation and retrieval reactivation (i.e., averaging retrieval-first and retrieval-second reactivation) were associated with greater integrated memory performance (*see Fig. S3 for statistical details*).

To investigate different emotion-modulated effects on hippocampal encoding reactivation across the three groups, we conducted a one-way ANOVA with Group (NN vs. NE vs. EE) as a between-subject factor (**Fig. 2C**). This analysis revealed a significant main effect of Group (F_(2,67)_ = 3.94, *p* = 0.024, partial ^2^ = 0.11). Post-hoc comparisons revealed greater encoding reactivation in NE (*p* = 0.010, 95%CI = [0.011, 0.077]) and EE (*p* = 0.046, 95%CI = [0.000, 0.064]) groups than the NN group. This result indicates that emotional information generally increases trial-specific reactivation in the hippocampus not only for related neutral memory in the NE group, but also for the other emotional memory in the EE group during encoding.

Subsequently, to investigate different emotion-modulated effects on hippocampal retrieval reactivation, we conducted a 2 (Measure: retrieval-first reactivation vs. retrieval-second reactivation) by 3 (Group: NN vs. NE vs. EE) repeated-measures ANOVA. This analysis revealed a significant Measure-by-Group interaction effect (F_(2,65)_ = 3.42, *p* = 0.039, partial η^2^ = 0.10; *Fig. S4*). Post-hoc comparisons revealed both greater retrieval-first reactivation (*p* = 0.019, 95%CI = [0.003, 0.027]) and retrieval-second reactivation (*p* = 0.033, 95%CI = [0.001, 0.026]) in the NE group than the NN group. However, it only revealed greater retrieval-second reactivation (*p* = 0.008, 95%CI = [0.005, 0.029]), but not retrieval-first reactivation (*p* = 0.39), in the EE group than the NN group. Furthermore, although only first associations were instructed for retrieval during the category recall test, the retrieval-second reactivation was significantly stronger than retrieval-first reactivation in the EE group (t_(21)_ = 2.91, *p* = 0.008, Cohen’s *d* = 0.62), but not in NN (t_(24)_ = −0.22, *p* = 0.828) and NE (t_(20)_ = −0.72, *p* = 0.483) groups. These results suggest that the hippocampal reactivations of first and second associations during retrieval might facilitate each other in the NE group, but interfere with each other in the EE group. To directly investigate this special interference effect between the retrieval of first and second associations in the EE group, we measured retrieval reactivation interference as a difference between retrieval-second and retrieval-first reactivation (i.e., retrieval-second minus retrieval-first reactivation). One-way ANOVA revealed a significant main effect of Group (F_(2,65)_ = 3.43, *p* = 0.039, partial η^2^ = 0.10) (**Fig. 2D**). Post-hoc comparisons revealed greater retrieval reactivation interference in the EE group than NN (*p* = 0.032, 95%CI = [0.001, 0.022]) and NE (*p* = 0.022, 95%CI = [0.002, 0.024]) groups. This result further indicates that emotional information impairs trial-specifc reactivation in the hippocampus for other related emotional memory in the EE group during retrieval.

To examine the hippocampal regional specificity for above observed effects, we conducted parallel control analyses for the posterior medial cortex (PMC). The PMC is known as a critical region engaged in retrieval-mediated reinstatement of initial encoding patterns (Chen et al., 2017; Jonker et al., 2018; Ranganath and Ritchey, 2012). However, these PMC analyses revealed no reliable effect for either encoding reactivation (F_(2,67)_ = 0.69, *p* = 0.503) or retrieval reactivation interference (F_(2,65)_ = 1.13, *p* = 0.330; *Fig. S5*).

Altogether, these reactivation results demonstrate that the emotion-related facilitation effect is associated with increased hippocampal reactivation during both encoding and retrieval. However, the emotion-related interference effect is associated with a potential trade-off between impaired hippocampal reactivation during retrieval and increased hippocampal reactivation during encoding. Both effects are selectively linked to hippocampal-mediated memory integration mechanisms, but not other regions involved in reactivation (or reinstatement), such as the PMC.

### Functional connectivity: The emotion-related facilitation and interference effects are associated with different strength levels of hippocampal connectivity during encoding

To further investigate whether and how hippocampal functional connectivity contributes to the emotion-related facilitation and interference effects, we conducted a task-dependent gPPI analysis with the hippocampal ROI as a seed (**Fig. 3A**). We focused on the processes of successful memory integration. Therefore, the hippocampal connectivity analysis was conducted only for the triplets with both associations remembered during the encoding of second associations relative to first associations. We extracted hippocampal connectivity data from the anatomically defined amygdala ROI (**Fig. 3B**) and then submitted the data into a one-way ANOVA. This analysis revealed a significant main effect of Group (F_(2,58)_ = 4.04, *p* = 0.023, partial η^2^ = 0.12; **Fig. 3D**). Post-hoc comparisons revealed greater hippocampal-amygdala connectivity in the NE group than NN (*p* = 0.021, 95%CI = [0.085, 1.006]) and EE (*p* = 0.013, 95%CI = [0.125, 1.035]) groups. Beyond the hippocampal-amygdala connectivity analysis, the whole-brain hippocampal connectivity analysis revealed significant main effects of Group in the right temporoparietal junction (TPJ), bilateral supplementary motor area (SMA), bilateral middle cingulate cortex (MCC) and left precuneus (**Fig. 3C**; *Table S1*). Post-hoc comparisons revealed significantly greater hippocampal functional coupling with these regions in the NE group than NN (TPJ: *p* < 0.001, 95%CI = [0.962, 2.401]; SMA: *p* < 0.001, 95%CI = [0.944, 2.308]; MCC: *p* < 0.001, 95%CI = [0.966, 2.139]; precuneus: *p* < 0.001, 95%CI = [1.076, 2.367]; **Fig. 3E**). It also revealed greater hippocampal coupling in the EE group than the NN group (SMA: *p* = 0.023, 95%CI = [0.111, 1.459]; MCC: *p* = 0.059, 95%CI = [-0.021, 1.138]; precuneus: *p* = 0.012, 95%CI = [0.193, 1.469]), whereas weaker than the NE group (TPJ: *p* = 0.001, 95%CI = [−1.948, −0.526]; SMA: *p* = 0.015, 95%CI = [−1.515, −0.167]; MCC: *p* = 0.001, 95%CI = [−1.574, −0.415]; precuneus: *p* = 0.007, 95%CI = [−1.528, −0.253]; **Fig. 3E**). These results indicate that emotional information generally enhances hippocampal coupling with the amygdala, TPJ, SMA, MCC, and precuneus supporting successful integrative encoding in the NE and EE groups, compared to the NN group. However, the emotional enhancement effect for hippocampal coupling in the EE group is weaker than it in the NE group. Additionally, no main effect of Group was found in the hippocampal connectivity analysis during retrieval (*Fig. S6*).

Together, these connectivity results demonstrate that the emotion-related facilitation effect is associated with strengthened hippocampal connectivity, but the interference effect is associated with relatively weak hippocampal connectivity during online integration.

## Discussion

In this study, we examined distinct neural mechanisms of different emotional modulations on memory integration. Specifically, we found that emotional information facilitated memory integration with related neutral information, but disrupted the integration with other emotional information. Emotion-facilitated memory integration of neutral and emotional information was associated with increased hippocampal reactivation during both encoding and retrieval. This facilitated integration was also associated with strengthened hippocampal connectivity with the amygdala, TPJ, SMA, MCC, and precuneus during integrative encoding. Emotion-interfered memory integration of two emotional pieces of information was associated with impaired hippocampal reactivation during retrieval, which seemed to offset the facilitating effect of increased hippocampal reactivation during encoding. This disrupted integration was associated with relatively weak hippocampal coupling with the SMA, MCC, and precuneus during integrative encoding. Our findings identify the emotion-induced facilitation and interference effects occurring on two types of emotional memory integration, through distinct dynamical hippocampal processes of trial-specific reactivation and connectivity.

### Emotion-facilitated memory integration

We provided compelling evidence for an emotion-induced facilitation effect on memory integration with neutral information. Behaviorally, the facilitation effect only occurred on the integrated memory in the immediate test. This result suggests a rapid and trial-specific facilitation for integrating related neutral and emotional memories, supporting the view of “associative facilitation” (Kuhl et al., 2010; Schlichting and Preston, 2014; Wimmer and Shohamy, 2012). It is also in line with previous studies, showing that linking a neutral mundane event to emotionally salient information can enhance memory for the neutral event due to its gained value and significance for the future (Holmes et al., 2022; Li et al., 2008; Sharpe et al., 2017; Zhu et al., 2022). However, we did not find any facilitation effect on non-integrated memories, which differs from the delayed and generalized emotional enhancement for weak memories encoded closely in time, as proposed in synaptic tagging-and-capture models (Ballarini et al., 2009; Dunsmoor et al., 2015; Frey and Morris, 1997). Thus, the observed emotion-facilitated effect appears to be specific for episode-unique integration, which maintains our emotional generalization to target events and avoids maladaptive overgeneralization resulting in affective disorders, such as phobia and posttraumatic stress disorder (PTSD) (Lange et al., 2019; Mary et al., 2020; Sripada et al., 2012).

In support of the emotion-facilitated integration, our imaging results showed that emotional information increased hippocampal reactivation of related memories during both encoding and retrieval. Consistent with emerging work using MVPA to study human episodic memory (Heinen et al., 2023; Hennings et al., 2022; Staresina et al., 2012; Xue et al., 2010), our findings of emotionally-charged increases in reactivation reflect trial-specific reinstatement (i.e., pair-specific similarity) controlling for a general category-level representation (i.e., across-pair similarity). Given the benefits of hippocampal pattern completion on integrative encoding (Kuhl et al., 2010; Shohamy and Wagner, 2008; Wong et al., 2019), the increased hippocampal reactivation of first neutral memories during encoding of second emotional associations could facilitate online integration by rapidly linking neural representations of related information. In addition, we found that the increased hippocampal reactivation of neutral memories during retrieval was accompanied by increased reactivation of non-targeted emotional memories. By the view of retrieval-induced facilitation among related memories (Anderson and McCulloch, 1999; Bäuml and Schlichting, 2014; Chan et al., 2006; Rowland and DeLosh, 2014; Wallner and Bäuml, 2017), the simultaneous reactivations of integrated neutral and emotional memories during retrieval could benefit each other, strengthen their connections and further facilitate their integration. Together, our findings demonstrate a dynamic emotion-facilitated integration mechanism, whereby emotional information firstly promotes the formation of integration during encoding and further strengthens this integration during retrieval, by increasing hippocampal reactivation of related neutral memories in both phases.

Moreover, we found strengthened hippocampal coupling with the amygdala, TPJ, SMA, MCC, and precuneus during encoding, contributing to this emotion-facilitated memory integration. The amygdala, with arousal-induced noradrenergic activation, is well recognized to play an essential role in emotional learning and benefit hippocampal-dependent memory. The SMA and MCC are also thought to regulate emotion generalization to target events along integrated memory traces but not to other irrelevant events (Kohn et al., 2014; Wager et al., 2008). The emotional involvement in memory processes promotes information transmission and communication between the hippocampus and neocortex (Battaglia et al., 2011; Hermans et al., 2014; Hofstetter et al., 2012; Zhu et al., 2022). Indeed, we found strengthened hippocampal connectivity with the TPJ and precuneus, which are core regions of the default mode network (DMN) (Hyatt et al., 2015; Schacter and Addis, 2007; Schacter et al., 2011). Recent studies have shown that the DMN is reliably engaged in ‘online’ processing (i.e., encoding), updating prior beliefs in light of new knowledge to simulate possible future (Yeshurun et al., 2021; Zadbood et al., 2022). Together, these findings reveal that the emotion-facilitated memory integration is linked to a hippocampal hub neural circuit. Specifically, emotional information might strengthen hippocampal-DMN interactions through enhanced hippocampal-amygdala/SMA/MCC coupling, which thereafter promote hippocampal-mediated reactivation and the integration of episodic memories.

### Emotion-interfered memory integration

While emotional information acts like a flashlight illuminating nearby representations, multiple emotional information can be dazzling and make it challenging to distinguish between these representations. We found evidence for an emotion-induced interference effect on memory integration with other emotional information. This interference effect is also selective for integrated memories, indicating its specificity for episode-unique integration. Additionally, we found that this interference effect might be due to a competitive relationship between two emotional memories. This result supports the “interference theory”, proposing that two salient emotional pieces of information interfere with each other because of their equal dominance in attention and memory (Itti and Koch, 2000; Mather, 2007, 2009). The mutual inhibition between related emotional information would lead to an overall suppression of their integration (Mather and Sutherland, 2011; Mitchell et al., 2006). This emotion-interfered integration is also adaptive, because it may protect our memory system from conflicts across various emotional information and intrusive traumatic recollections (Brewin, 2006; Wimber et al., 2008). It thereby relieves emotional overload and dysregulation with vulnerability to affective disorders, such as anxiety, depression, phobias, and PTSD (Papageorgiou et al., 2000; Wells et al., 2004; Wells et al., 1997).

Consistent with emotion-facilitated integration, we found increased hippocampal reactivation of first emotional memories during the encoding of second emotional associations in the emotion-interfered integration. Besides reactivation, we also found strengthened hippocampal connectivity with the SMA, MCC, and precuneus supporting successful integrative encoding. This pattern of results suggests that online processing is generally promoted in the both facilitated and interfered integrations with new emotional learning. It is worth noting that the promoting effect seems to be weaker in emotion-interfered integration, considering its relatively weak statistical effect of reactivation and hippocampal-neocortical interactions compared with the emotion-facilitated integration.

More importantly, our observed emotion-interfered integration was mainly contributed by impaired hippocampal reactivation during retrieval, due to the competitive retrieval of two related emotional memories. We found a significantly increased hippocampal reactivation of non-target second emotional associations when first emotional associations were retrieved. The retrieval reactivation of non-target memories was even stronger than target ones, indicating an involuntary intrusive effect of recent emotional stimuli (Gagnepain et al., 2017; Herz et al., 2020; Mary et al., 2020). The view of mutually inhibitory control mechanisms is proposed to solve the conflict between two equally prioritized emotional pieces of information, by overall suppressing emotional benefits on hippocampal reactivation and weakening connections between related emotional memories (Brewin, 2006; Mather and Sutherland, 2011; Wimber et al., 2015). Furthermore, the interference effect of impaired retrieval reactivation potentially offsets the facilitating effect of increased encoding reactivation, ending up with a disrupted integration. Together, our findings demonstrate a dynamic emotion-interfered integration mechanism, whereby the emotional memory integration formed and promoted during encoding is later disrupted by impaired hippocampal reactivation of related emotional memories during retrieval.

## Conclusion

Our study specifies the emotion-induced facilitation and interference effects on two typical types of emotional memory integration, involving distinct dynamical processes of hippocampal trial-specific reactivation and connectivity. The emotion-facilitated memory integration of neutral and emotional information, is formed with increased hippocampal reactivation and connectivity during encoding, and further facilitated with mutually beneficial hippocampal reactivation of both information during retrieval. The emotion-interfered memory integration of two emotional pieces of information, is also formed with increased hippocampal reactivation and relatively weakly strengthened hippocampal connectivity during encoding, but later suffers more interference due to mutually impaired hippocampal reactivation during retrieval. Our findings provide a comprehensive explanation, with respect to the valences of related information, for previously divergent studies of emotional modulation on memory integration. These findings advance our understanding of neurobiological mechanisms by which emotions can differently modulate memory integration to foster adaptation to the future, and also provide novel insights into emotion dysregulation and maladaptive generalization in mental disorders.

## CRediT author statement

**Yannan Zhu**: Conceptualization, Formal analysis, Data curation, Writing - original draft, Writing – review & editing, Visualization, Project administration, Funding acquisition; **Wei Liu**: Conceptualization, Formal analysis, Investigation, Data curation, Writing – review & editing, Project administration, Funding acquisition; **Nils Kohn**: Conceptualization, Data curation, Writing – review & editing, Supervision; **Guillén Fernández**: Conceptualization, Data curation, Writing – review & editing, Supervision, Project administration, Funding acquisition.

## Acknowledgments

This work was supported by the Open Research Fund of the State Key Laboratory of Cognitive Neuroscience and Learning (CNLYB2103, W.L.), the Open Research Fund of the Key Laboratory of Adolescent Cyber Psychology and Behavior (CCNUCYPSYLAB2022B10, W.L.), the Major Program of the National Social Science Foundation of China (22&ZD187, W.L.), and the Ph.D. fellowship of the Chinese Scholarship Council (201806040186, Y.Z.). We thank Nancy Peeters for the assistance of data acquisition, Merel Koning and Bas Meuter for the memory performance evaluation.

