## Supplemental Materials for "Emotional information facilitates or disrupts memory integration through distinct hippocampal processes of reactivation and connectivity"

### Methods

#### Participants

All participants were recruited from the Radboud Research Participation System (<https://radboud.sona-systems.com>). Participants were randomly assigned by an electronic data capture system (Castor, <https://www.castoredc.com/>) into one of the three groups, NN (n = 25; 14 females; mean age ± s.d., 23.8 ± 2.98 years), NE (n = 21; 16 females; mean age ± s.d., 23.0 ± 3.90 years) and EE (n = 24; 16 females; mean age ± s.d., 23.2 ± 2.95 years) groups. All participants had normal or corrected-to-normal vision, and reported no history of any neurological diseases or psychiatric disorders.

#### Imaging preprocessing

Brain imaging data were organized into the Brain Imaging Data Structure (BIDS) (Gorgolewski et al., 2016), and preprocessed using fMRIPrep (version 20.0.6) (Esteban et al., 2019) based on Nipype (version 1.4.2) (Gorgolewski et al., 2011) and FEAT (fMRI Expert Analysis Tool, version 6.0) as part of FSL (FMRIB’s Software Library, <http://www.fmrib.ox.ac.uk/fsl>) (Jenkinson et al., 2012).

#### Univariate GLM analysis

These stick functions were generated by FSL’s Motion Outliers algorithm, which identifies large displacements in head position by measuring the difference in intensity between each volume and the preceding volume.

#### Reactivation: MVPA

The stick function regressors in the above univariate GLMs were also included in each MVPA GLM to account for sudden head movements. We eliminated the mean activation (i.e., *t*-value) from each pattern by z-scoring across voxels within the ROI. Thus, the resulting mean-centered pattern has relative voxel amplitudes (i.e., voxel-level variability) preserved without any difference in the overall amplitudes, ensuring that the subsequent similarity results are fully attributable to the pattern itself (Coutanche, 2013; Tompary and Davachi, 2017).

#### Functional connectivity: PPI

The physiological activity of the given hippocampal seed region was computed as the mean time series of all voxels. They were then deconvolved to estimate neural activity (i.e., physiological variable), and multiplied with the task design vector by contrasting the encoding of second and first associations (or the retrieval of first associations in triplets with both associations remembered and triplets with no association remembered) (i.e., psychological variable) to form a psycho-physiological interaction vector. This interaction vector was convolved with FEAT’s HRF to form the PPI regressor of interest. Another psychological variable representing the task conditions (i.e., the encoding of second plus first associations, or the retrieval of first associations in triplets with both associations remembered plus triplets with no associations remembered) was also included in the GLM to remove out the effects of common driving inputs on brain connectivity. The stick function regressors in the above univariate GLMs were also included in each PPI GLM to account for sudden head movements.

##### **Table S1. Brain regions involved in successful integration effect of hippocampal connectivity during encoding**

| **Brain Regions** | **Hemisphere** | ***Z* values** | **MNI Coordinates** | | |
| --- | --- | --- | --- | --- | --- |
|  |  |  | **X** | **Y** | **Z** |
| **Second associations vs. First associations** | | | | | |
| Inferior parietal lobule | R | 4.68 | 56 | -26 | 26 |
| Supplementary motor area | R | 5.07 | 4 | -14 | 58 |
| Middle cingulate cortex | R | 4.83 | 2 | -6 | 44 |
| Precuneus | L | 4.58 | -24 | -42 | 72 |
| Precentral Gyrus | R | 4.31 | 20 | -30 | 68 |
| Postcentral Gyrus | R | 4.30 | 12 | -56 | 72 |

Notes: Regions were derived from the hippocampal functional connectivity analysis for the triplets with both associations remembered during the encoding phase. Significant clusters, at voxel-wise z > 3.1 and cluster-level *p* < 0.05 with family-wise error correction for multiple comparisons, are reported with local maximum *Z* statistic in Montreal Neurological Institute (MNI) space. L, left; R, right.

**Figure S1**

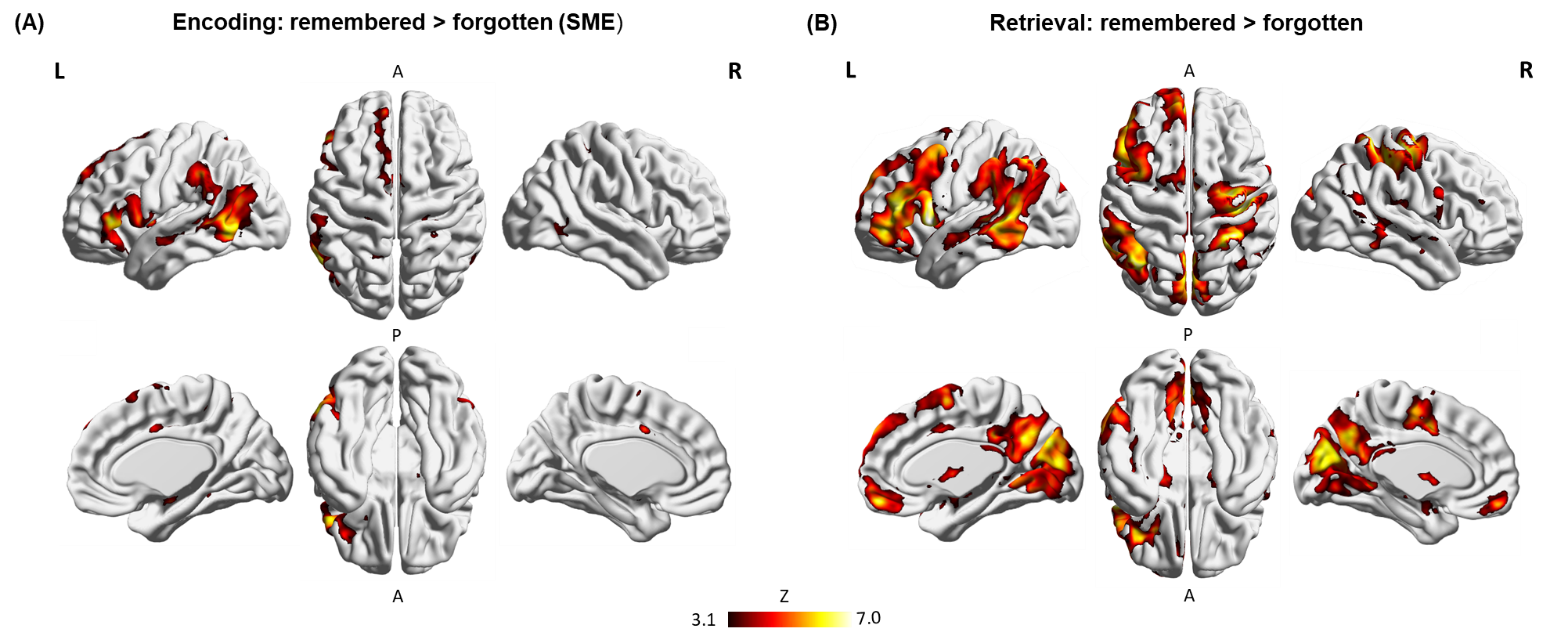

#### **Figure S1. Brain systems involved in subsequent memory effect during encoding and successful retrieval effect during retrieval.**

**(A)** Widespread brain activation associated with a subsequent memory effect (i.e., remembered vs. forgotten) during encoding. **(B)** Widespread brain activation associated with a successful retrieval effect (i.e., remembered vs. forgotten) during retrieval. Notes: Color bar represents z values. L, left; R, right; A, anterior; P, posterior.

**Figure S2**

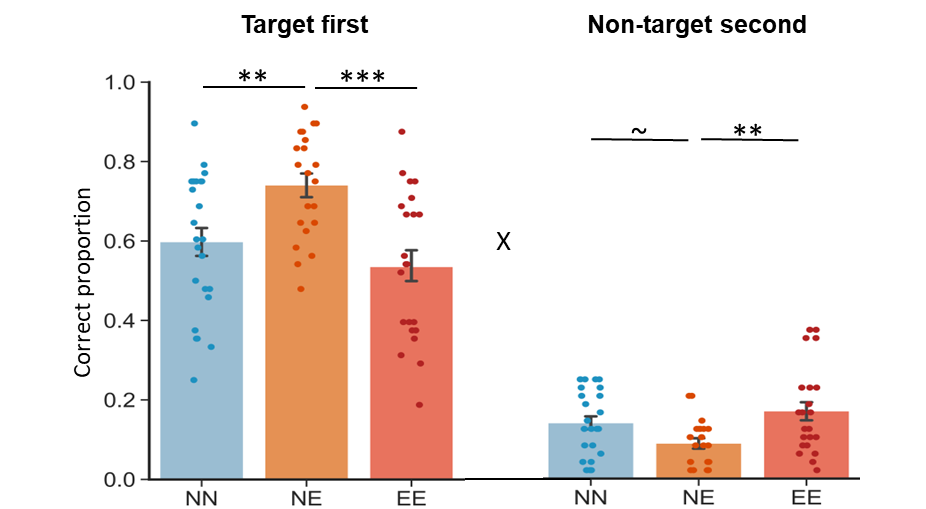

#### **Figure S2. Category-recall memory performance.** Bar graphs depict average correct proprotions for target first and non-target second associations in the three groups separately. Besides a significant interaction effect, the 2 (Memory: target first vs. non-target second) by 3 (Group: NN vs. NE vs. EE) repeated-measures ANCOVA (gender as a covariate of no interest) revealed significant main effects of Memory (F_(1,64)_ = 14.78, *p* < 0.001, partial η^2^ = 0.19) and Group (F_(2,64)_ = 5.93, *p* = 0.004, partial η^2^ = 0.16). Error bars represent the standard error of the mean. Dots represent data from each participant. “X” indicates significant interaction (*p* < 0.05). Notes: ~ *p* < 0.085; ***p* < 0.01; ****p* < 0.001; two-tailed tests.

**Figure S3**

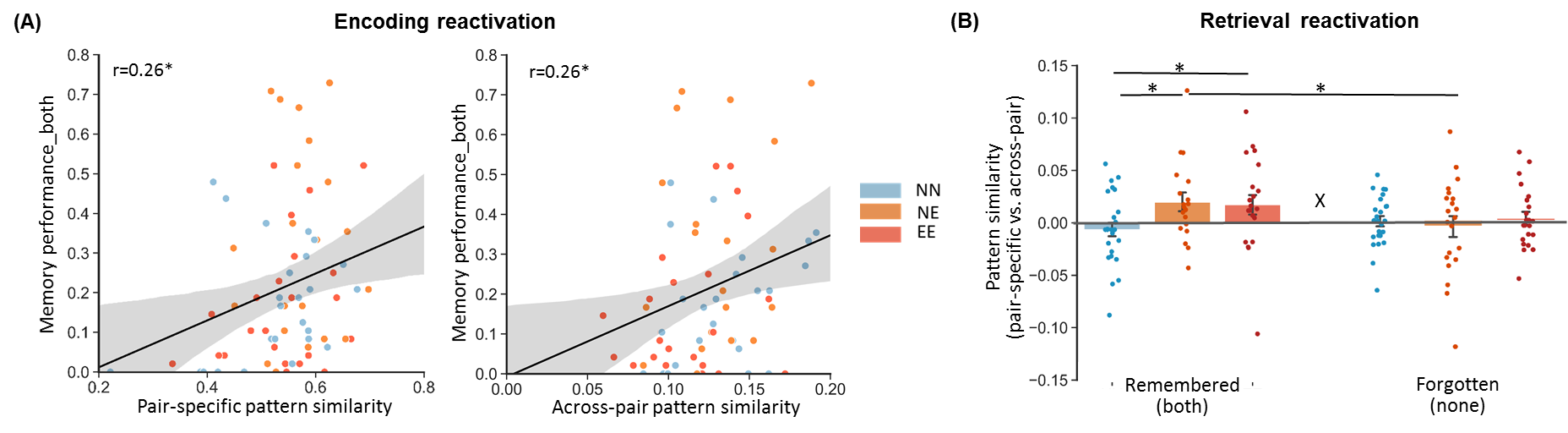

#### **Figure S3. Relationships between reactivation and memory performance.**

**(A)** Scatter plots depict correlations (i.e., a bootstrapped Pearson’s correlation analysis with 1000 samples) of integrated memory performance (i.e., memory for triplets with both associations remembered) with pair-specific (*left;* r = 0.26, *p* = 0.032, 95%CI = [0.058, 0.441]) and across-pair (*right;* r = 0.26, *p* = 0.028, 95%CI = [0.053, 0.471]) pattern similarities during encoding. Solid lines indicate the best linear fit, and light shadows indicate 95% confidence intervals. **(B)** Bar graphs depict average retrieval reactivation (i.e., averaging retrieval-first and retrieval-second reactivation) for remembered (i.e., both associations remembered) and forgotten (i.e., no association remembered) triplets in the three groups separately. A 2 (Memory: remembered vs. forgotten) by 3 (Group: NN vs. NE vs. EE) repeated-measures ANOVA revealed a significant main effect of Memory (F_(1,62)_ = 5.29, *p* = 0.025, partial η^2^ = 0.08), with greater retrieval reactivation for remembered than forgotten triplets. Error bars represent the standard error of the mean. Dots represent data from each participant. “X” indicates significant interaction (*p* < 0.05). Notes: **p* < 0.05; two-tailed tests.

**Figure S4**

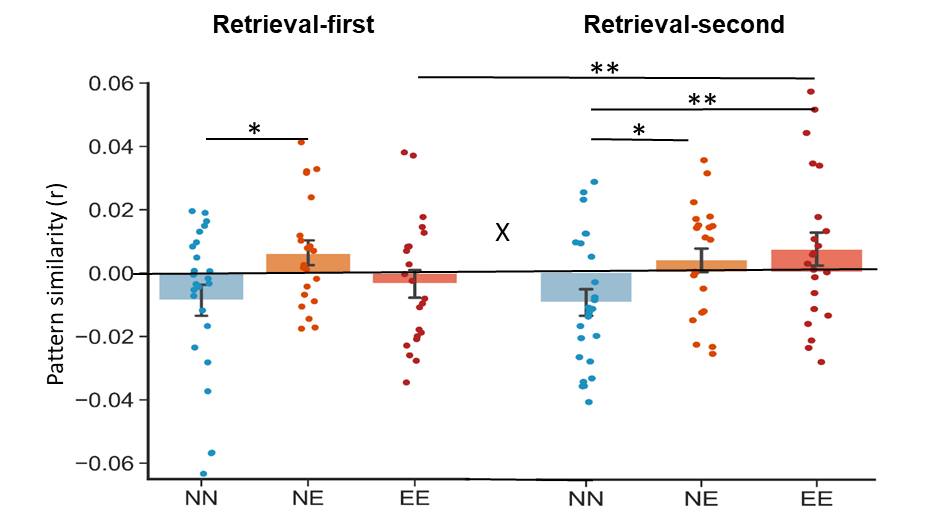

#### **Figure S4. Retrieval reactivation of first and second associations.**

Bar graphs depict average retrieval-first reactivation and retrieval-second reactivation (i.e., retrieval-first/second pair-specific relative to retrieval across-pair similarity) in the three groups separately. Error bars represent the standard error of the mean. Dots represent data from each participant. “X” indicates significant interaction (*p* < 0.05). Notes: **p* < 0.05; ***p* < 0.01; two-tailed tests.

**Figure S5**

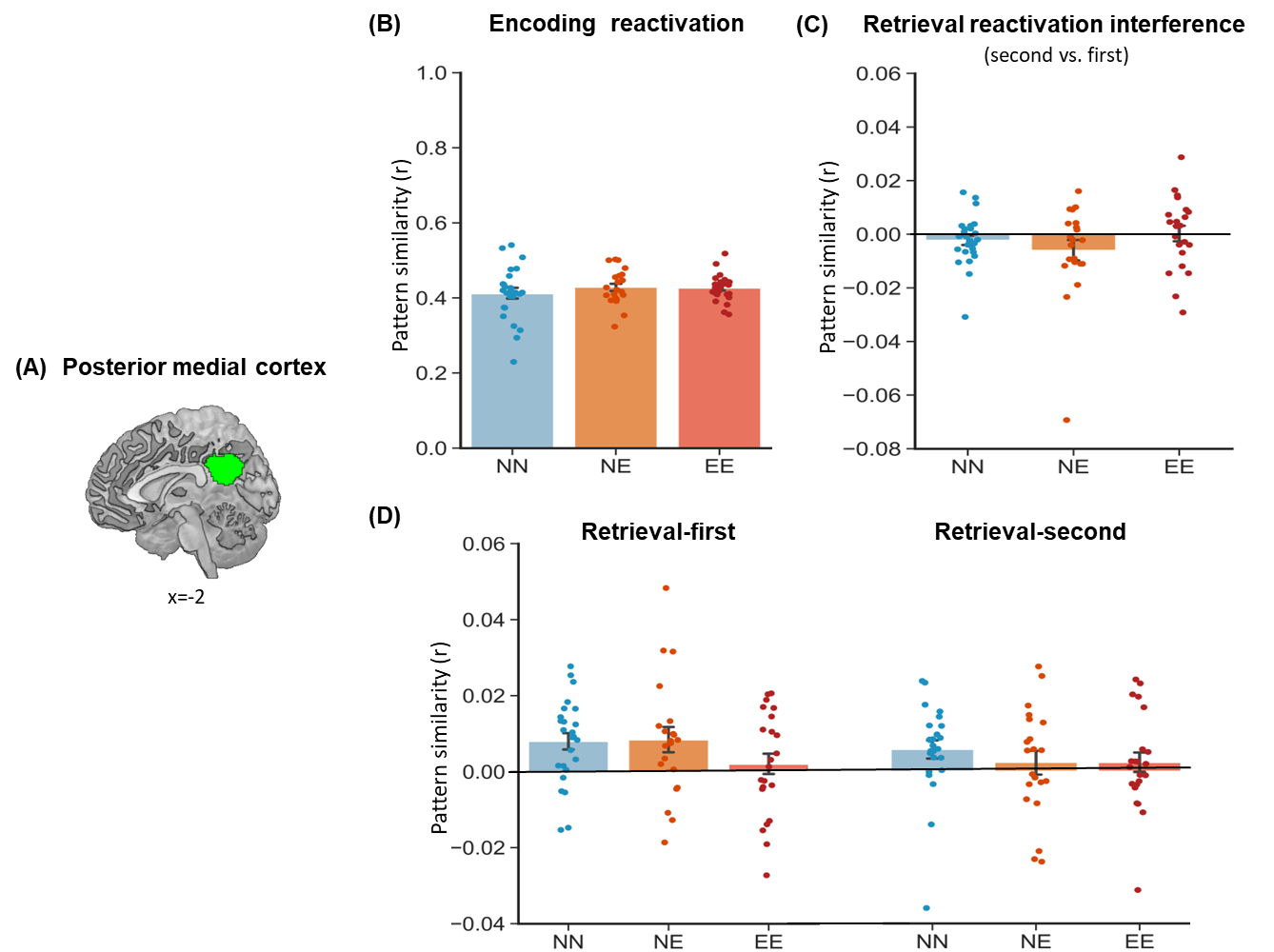

#### **Figure S5. PMC reactivation during encoding and retrieval.**

**(A)** The PMC ROI was used in parallel control pattern similarity analyses. **(B-D)** Bar graphs depict average encoding reactivation, retrieval reactivation interference and retrieval-first/second reactivation in the three groups separately. Error bars represent the standard error of the mean. Dots represent data from each participant.

**Figure S6**

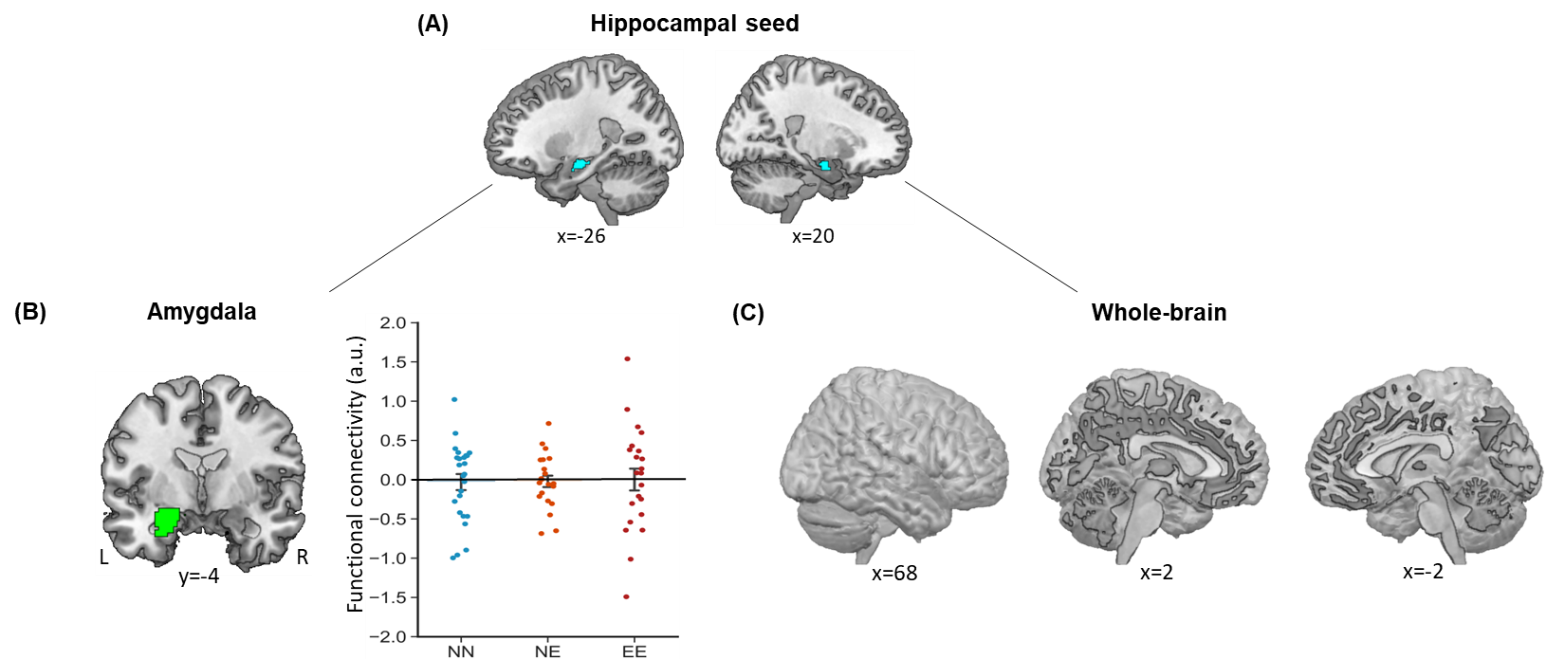

#### **Figure S6. Hippocampal connectivity involved in retrieval-mediated memory integration.**

**(A)** The bilateral hippocampal seed was used in task-dependent gPPI analysis (i.e., the retrieval of first associations in triplets with both associations remembered vs. triplets with no association remembered). **(B)** The anatomical amygdala ROI was used in the hippocampal connectivity analysis. Bar graphs depict average hippocampal-amygdala connectivity during retrieval in the three groups separately. The main effect of Group was non-significant (F_(2,65)_ = 0.03, *p* = 0.970). **(C)** No significant cluster, surviving at the thresholding criteria used in the present study (i.e., voxel-wise z > 3.1, cluster-level *p* < 0.05 FWER corrected), showed the main effect of Group during retrieval. Error bars represent the standard error of the mean. Dots represent data from each participant. Notes: a.u., arbitrary unit; L, left; R, right.
